# Unravelling intra- and intersegmental neuronal connectivity between central pattern generating networks in a multi-legged locomotor system

**DOI:** 10.1101/453423

**Authors:** Silvia Daun, Charalampos Mantziaris, Tibor I. Tóth, Ansgar Büschges, Nils Rosjat

## Abstract

Animal walking results from a complex interplay of central pattern generating networks (CPGs), local sensory signals expressing position, velocity and forces generated in the legs, and coordinating signals between neighboring ones. In the stick insect intra- and intersegmental coordination is conveyed by these sensory signals. The CPGs control the activity of motoneuron pools and are thereby responsible for the generation of rhythmic leg movements. The rhythmic activity of the CPGs can be modified by the aforementioned sensory signals. However, the precise nature of the interaction between the CPGs and these sensory signals has remained largely unknown. Experimental methods aiming at finding out details of these interactions often apply the muscarinic acetylcholine receptor agonist, pilocarpine in order to induce rhythmic activity in the CPGs. Using this general approach, we removed the influence of sensory signals and investigated the putative connections between CPGs associated with the coxa-trochanter (CTr)-joint in the different segments (legs) in more detail. The experimental data underwent connectivity analysis using Dynamic Causal Modelling (DCM). This method can uncover the underlying coupling structure and strength between pairs of segmental ganglia (CPGs). For the analysis we set up different coupling schemes (models) for DCM and compared them using Bayesian Model Selection (BMS). Models with contralateral connections in each segment and ipsilateral connections on both sides, as well as the coupling from the meta-to the ipsilateral prothoracic ganglion were preferred by BMS to all other types of models tested. Moreover, the intrasegmental coupling strength in the mesothoracic ganglion was the strongest and most stable in all three ganglia.

## Introduction

Various experiments on vertebrates and invertebrates confirm the existence of central pattern generating networks (CPGs). These networks are responsible for the generation of periodic muscle activity in a given leg [14, 29]. The movement of each leg has to be coordinated with that of the other legs in order to produce walking. In the stick insect *Carausius morosus*, each leg is individually controlled by its own CPGs located in the pro- (front legs), meso- (middle legs) and metathoracic ganglion (hind legs) [15, 46]. Each leg consists of three main leg joints about which leg segments execute coordinated movements during walking and climbing. The thorax-coxa (ThC) joint is responsible for forward and backward movements, the coxa-trochanter (CTr) joint enables the femur to move in upward and downward direction. The femur-tibia (FTi) joint brings about flexing and stretching of the leg by moving the tibia relative to the femur. Each of the leg joints is associated with an antagonistic muscle pair: the protractor-retractor (ThC), the levator-depressor (CTr) and the flexor-extensor (FTi) muscle pair [19]. The rhythmic (periodic) activation of the muscles originates in the corresponding CPGs [5] and sensory signals play a significant role in coordinating the motor output between segments [2–4, 21, 25]. In addition, a previous study showed that, in the absence of sensory feedback, depressor CPG activity is weakly coupled to the one of all other segments [28]. The authors demonstrated that the different intrasegmental phase relationships for isolated ganglia were stabilized in the case of interconnected ganglia.

However, little is known about the interaction, i.e. the strength and the nature of the couplings between the different CPG networks. In order to understand how a stable locomotor pattern is generated, we need to understand the contribution of the central and peripheral sensory signals, and the interactions between them. Up to now, there has been a long and successful history of mathematically describing and modelling central pattern generating networks by means of phase oscillators [10]. A variety of interesting and fruitful insights have been gained from the models of weakly coupled oscillators [9, 27, 37]. Nevertheless, so far no method has been found to estimate the coupling strengths *from the data* and not from the model itself. This means that in the stick insect in particular, the connection strengths between the various CPGs in the different ganglia have not yet been studied (to the best of our knowledge also not for other insects at least not for the whole nerve cord).

There are some promising approaches concerning electroencephalography (EEG) and magnetoencephalography (MEG) recordings, e.g. [23, 33, 35]. In this paper, we tackle the problem of estimating the coupling architecture and strengths by using a modelling approach called Dynamic Causal Modelling (DCM). It serves the purpose of assigning relative coupling strengths to the (neuronal) connections between the levator-depressor CPGs in different hemisegments. In contrast to other methods, e.g. phase-coupling, coherence or weakly coupled oscillators, this approach has three major advantages. First, DCM is able to give insight into effective, i.e. directed, connectivity while many other approaches only focus on functional, i.e. undirected, connectivity. Second, DCM allows to use Bayesian Model Selection (BMS) to determine the model architecture best fitting the data. And third, in contrast to other approaches it is also applicable in the context of few experimental recordings, as long as these are of a sufficient length.

To validate the estimates on the strength of the coupling between the CPGs of the different segments in the thoracic nerve cord of the stick insect obtained with this modeling approach, a descriptive data analysis method was used.

This article is organized as follows: in section 1, *Materials and methods*, we will review the experimental methods and explain the basic properties of the Dynamic Causal Modelling (DCM) including Bayesian Model Selection (BMS) as well as the descriptive phase-connectivity (PC) approach, which we use for the data analysis in the present work. Section 2, *Results*, is divided into three parts. First, we present the results of the DCM analysis for the meso- and metathoracic ganglia, then, the ones for the pro- and mesothoracic ganglia. We validate these results using the PC approach as well as additional experiments with known connectivity. Finally, we present the coupling architecture and strengths for the whole thoracic nerve cord, i.e. the pro-, meso- and metathoracic ganglia hypothesized by DCM. After all, the sections 3, *Discussion*, and 4, *Conclusion*, follow.

## 1 Materials and methods

### Animals

The experiments were carried out on adult female Indian stick insects of the species *Carausius morosus* [45]. The animals are obtained from the colony at the University of Cologne maintained at 22-24° C, at approximately 60% humidity and under a 12 h light / 12 h dark cycle.

### Preparation

We extracellularly recorded the rhythmic activity from C2 leg nerves, which contain the axons that innervate the slow and the fast *depressor trochanteris* muscles (SDTr and FDTr respectively) [39]. We did so with the nerves in the contralateral pro-, meso- and metathoracic ganglia using ‘hook’ electrodes [40] in an isolated and deafferented preparation. To this end, all legs of the stick insects were removed, all lateral and connective nerves, except the ones of interest, were cut off. Also axons of sensory neurons were destroyed to prevent sensory feedback and peripheral input from being recorded. Rhythmic activity in leg motoneuron (MN) pools was then induced by bath application of 5 – 7 mM of the muscarinic receptor agonist *pilocarpine* [5]. For a detailed description of the preparation, experimental setup and electrophysiology see [28]. Fig 1 shows sample recordings from meso-meta, pro-meso and pro-meso-meta recordings.

**Fig 1.**
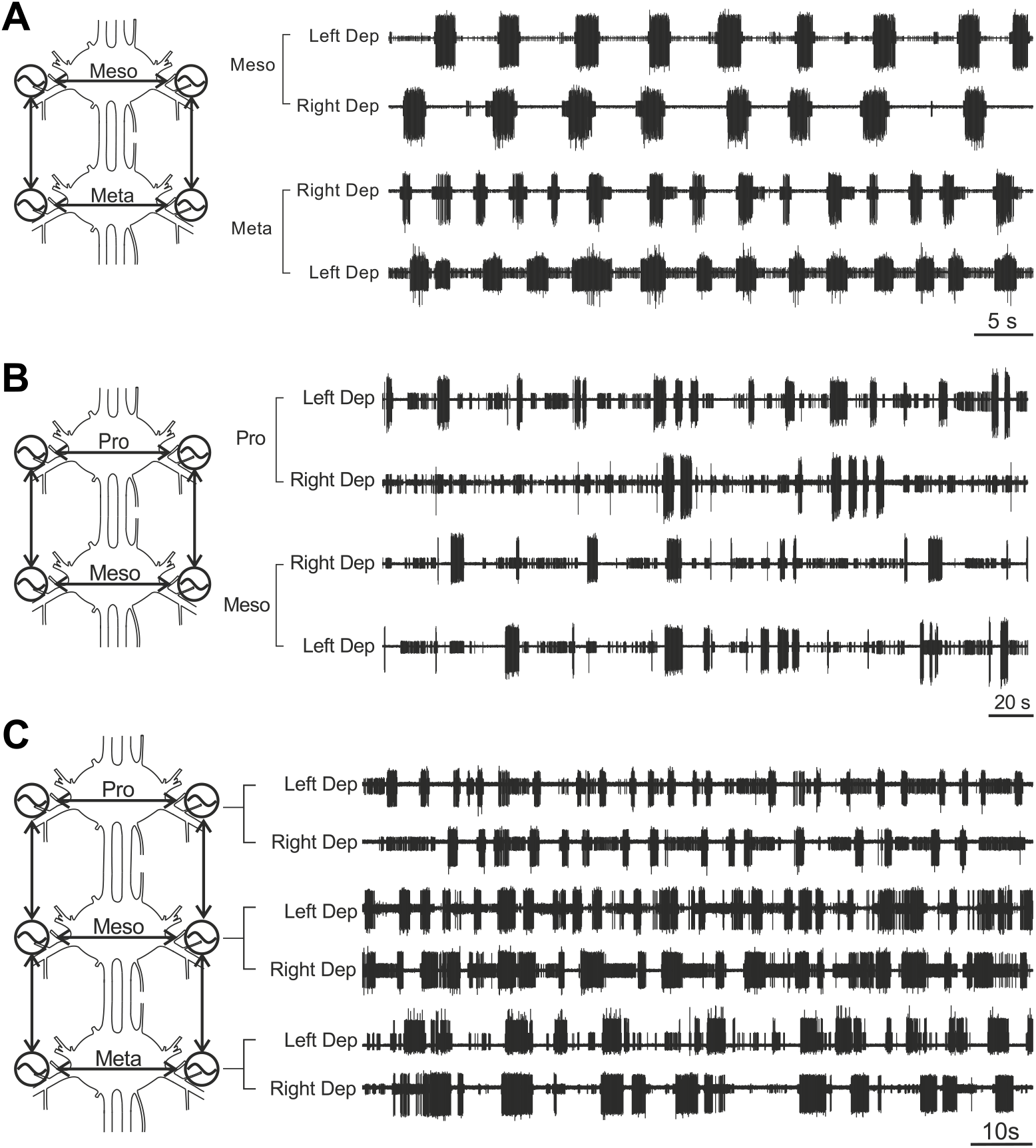
Sample recordings. Examplary recordings from left and right depressor nerves from A) 5 s of Meso-Meta recordings (218.72 *s* < *t* < 783.21 *s, N* = 10), B) 20s of Pro-Meso recordings (633.84 *s* < *t* < 2061.39 *s, N* = 13) and C) 10 s of Pro-Meso-Meta recordings (390.03 *s* < *t* < 790.97 *s, N* = 3).

### Data preprocessing

The collected data were preprocessed offline using Spike2 7.09 (CED, Cambridge, UK). First, we used the signal processing functions DC-remove, Rectify and Smooth to get rectified and smoothed waveforms that are corrected for DC (direct current) shifts (i.e., having a mean amplitude of zero). These data were then downsampled to 200 Hz and extracted as a time-series.

### Connectivity analysis

The data was further processed with MatLab R2011b (The MathWorks Inc., Massachusetts, USA) and Python 2.7.14. For Dynamic Causal Modelling (DCM), we used the Statistical Parametric Mapping toolbox (SPM12, Wellcome Trust Centre for Neuroimaging, London, UK) implemented in MatLab. Phase-difference analysis was done using custom programmed MatLab scripts and clustering algorithms implemented in the Python toolbox sci-kit learn [34].

### Dynamic Causal Modelling approach

We made use of Dynamic Causal Modelling (DCM) [23] to investigate the type and strength of intra- and intersegmental coupling between the thoracic ganglia of the levator-depressor system of the stick insect. This approach was developed for data recorded from the human brain and is widely used in the analysis of couplings in M/EEG and fMRI data [6, 20, 44], and in the analysis of local field potentials [33].

DCM uses neural mass models [31, 32] to describe the neuronal activity of the recorded sources (in our case CPGs). As there is no stimulation in our experimental setup, *i.e. absence of sensory input*, we selected the DCM for cross-spectral density (CSD) approach that does not include any inputs and is suitable for modelling steady-state like data [16]. This particular DCM approach is based on a linearization of dynamical systems, i.e. neuronal subpopulations are coupled via their mean fields. The connectivity is determined by modeling coherence and phase-differences of the observed electrophysiological measurements. A model is optimized using empirical measures of cross-spectral densities. In particular, the parameter values are fitted within a system of differential equations according to a predefined coupling structure (see Statistical Methods) to model, i.e. explain, the recorded source activity:

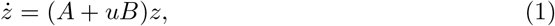

where z is the output activity of the sources. The coupling structure is defined in the matrix A, and possible changes in connectivity between different experimental conditions are modelled by using B. The coupling strengths saved in A and B are then fitted in order that *z*(*t*) has, in some well-defined sense, the smallest distance from the recorded activity, i.e. being optimal in that sense.

The connectivity strengths obtained from DCM underwent two independent validations: (i) by a phase-connectivity approach (see below) and (ii) through the analysis of an experimental condition with known connectivity, i.e. cut connectives between the segmental ganglia.

### Phase-connectivity approach (PC)

We validated the DCM connectivity results by analyzing, *in the absence of sensory input*, the mutual relationship of the rhythmic motor activities in the stick insect in order to uncover possible phase-coupling between them. The analysis was performed by means of established methods that are described elsewhere [37, 43]. Using this approach, we gained information about the time evolution of multiple rhythms propagating intra- or intersegmentally. The intersegmental analysis was done for the meso- and metathoracic ganglia first and then for the pro- and mesothoracic ganglia. We analyzed the phase of the rectified and smoothed signal obtained from each nerve as the activity evolved in time. For automatic and objective detection of burst onsets the preprocessed extracellular recordings were used to construct a discrete-time analytic signal

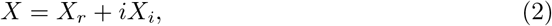

in the complex plane. Here, *X_r_* is the real data vector, and *X_i_* is the Hilbert transform of *X_r_* [30]. Then a Poincare-section was used to define the onsets of the bursts and thereby the reference phase of the rhythm. To obtain information on the phase *φ* of each recording, we linearly interpolated the phase angle between each pair of onsets of each single nerve and normalized it to lie in the interval [0,1) (mod 2*π*) during one cycle. As a last step, the phase was unwrapped, i.e. it grew monotonically, as if it were an ‘ordinary’ non periodic time signal (Fig 2, A left). When recording from the prothoracic ganglion, the signals showed activities with small amplitudes, in addition to the large-amplitude bursts. Thus, we modified the analysis by adjusting the Poincare-section such that only the big amplitudes were marked as burst onsets (Fig 2, A right; second black circle in the top panel). The adjustment was based on the k-means clustering algorithm implemented in the sci-kit learn toolbox in Python [26, 34] for k=2, which is motivated by separating the bursts into two clusters referring to the small and large units observed in the recordings. This method optimizes two centroids representing the small and big amplitude peaks and then assigns each detected peak to the closest cluster.

**Fig 2.**
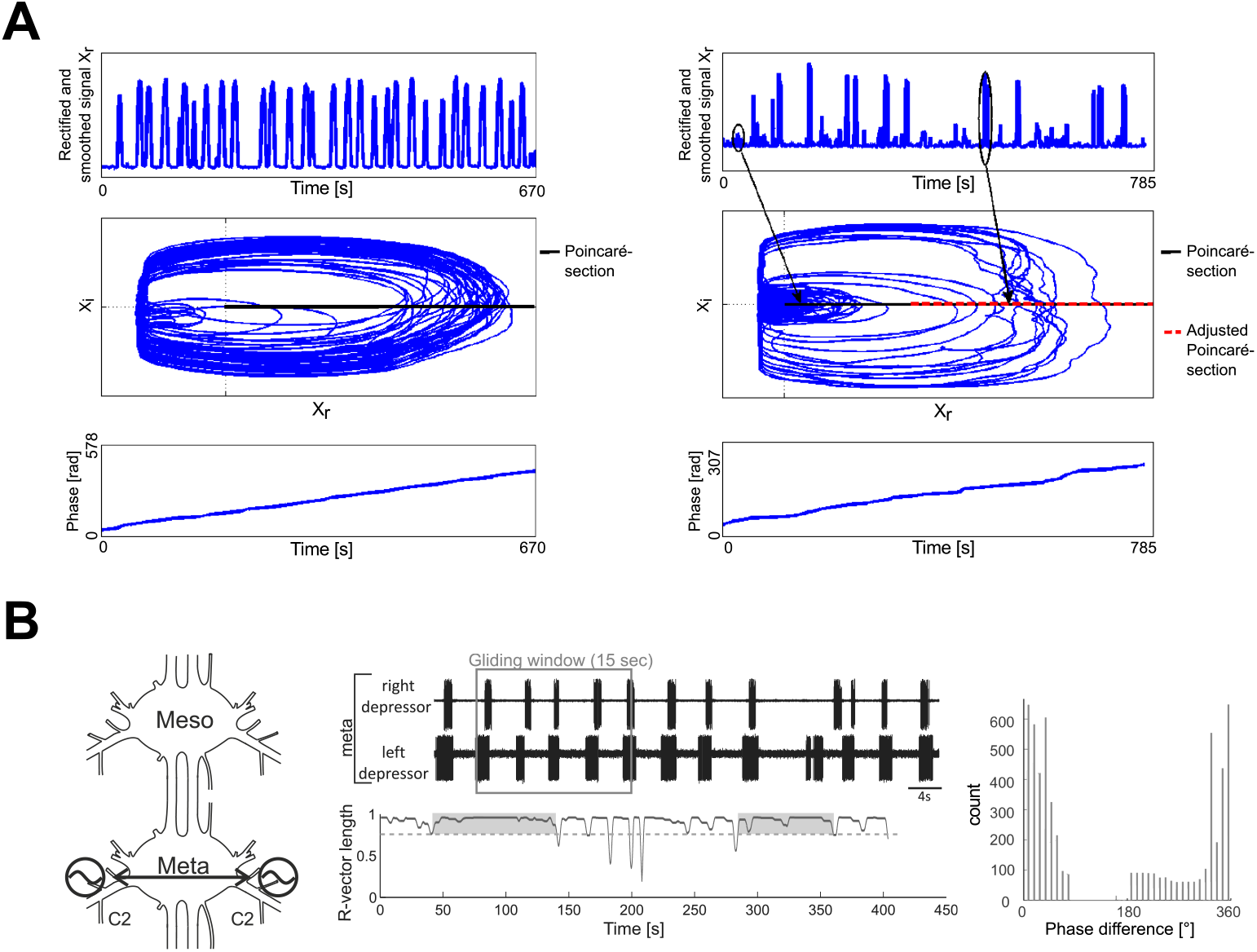
Phase-connectivity approach. A: Top - Rectified and smoothed signal *X_r_* (Left: meso-meta, Right: pro-meso); Middle - Discrete-time analytic signal with Poincare-section marked in black (normal) and red (eliminating small units) (Left: meso-meta, Right: pro-meso); Bottom - Resulting instantaneous (unwrapped) phase *φ* (Left: meso-meta, Right: pro-meso). B: Left - Schematic drawing which shows which segmental depressor activity was recorded and analyzed. Middle top - Sample recording of left and right depressor activity in the metathoracic ganglion; Middle bottom - Time evolution of the R-vector length of left-right metathoracic phase-differences over the time span of a recording, intervals in which coupling occurs are marked with grey boxes; Right - Phase-histogram shown for the coupled interval ≈ [50,140] (see text for details).

To investigate the coupling of two CPGs, we calculated their phase-difference. The signals are considered to be coupled if their phase-difference remains constant over a longer time-period, i.e.

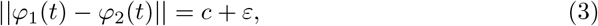

*ε* being a small error (compared to *c*). In our analysis, we ensured this by requiring the two criteria below to be fulfilled for the R-vector *R* (cf. [1]), which is defined by

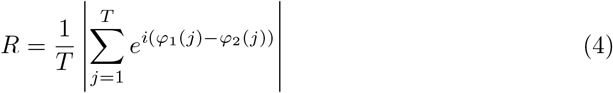

with the number of time points *T* and the corresponding phase-difference *φ*_1_(*j*) – *φ*_2_(*j*). It measures the similarity of the phase-differences of the coupled phases and ranges from 0 (random phase-differences) to 1 (identical phase-differences):

1. The R-vector in a 15 s long gliding time-window should be larger than 0.8 for at least 50 s (corresponding to 10 cycles) (Fig 2, B middle bottom panel).
2. Over the whole time-interval defined in 1., there should be a clear peak in the histogram (Fig 2, B right), with an R-vector greater than 0.3. This criterion prevents drifting of the phases over the interval where coupling exists.

Both thresholds for the R-vector lengths were adjusted manually in such a way that the program was able to correctly assign clearly coupled or clearly uncoupled intervals to the correct group.

After phase-coupled intervals were identified, we defined the strength of the coupling to be the likelihood of coupling over the whole recording, i.e. the sum of all interval lengths in which coupling occurred, divided by the total length of the corresponding recording. Recordings with no coupling were taken into account with 0 s of coupling.

### Statistical Methods

The coupling architecture best fitting the data in the DCM approach is determined by Bayesian Model Selection (BMS) [36]. This method yields two measures: (a) the relative logarithmic model evidence and (b) the model posterior probability. The logarithmic model evidence, log *B_ji_*, of a model *j* is displayed relative to the least probable model *i*

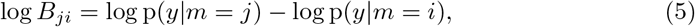

with the probability p(*y|m* = *j*) describing the likelihood of the observed data being generated by model j. Given equal priors *p*(*m* = *i*) = *p*(*m* = *j*) (for the different models) the posterior probability of the *i*-th model *p*(*m* = *i|y*) is

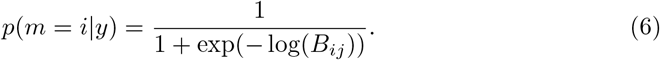

Finding the best coupling structure via BMS enables us to investigate intra- and intersegmental coupling strengths in the preferred model architecture. For the meso-meta and pro-meso networks, we tested for 7 different network architectures by stepwise reducing the number of connections from fully connected to fully unconnected (see Fig 3, A). In the case of pro-meso-meta connectivity we based our models on the results of the meso-meta and pro-meso analyses and added possible, physiologically as well as theoretically motivated, pro-meta connections to the coupling structure (see Results and Fig 3, B).

**Fig 3.**
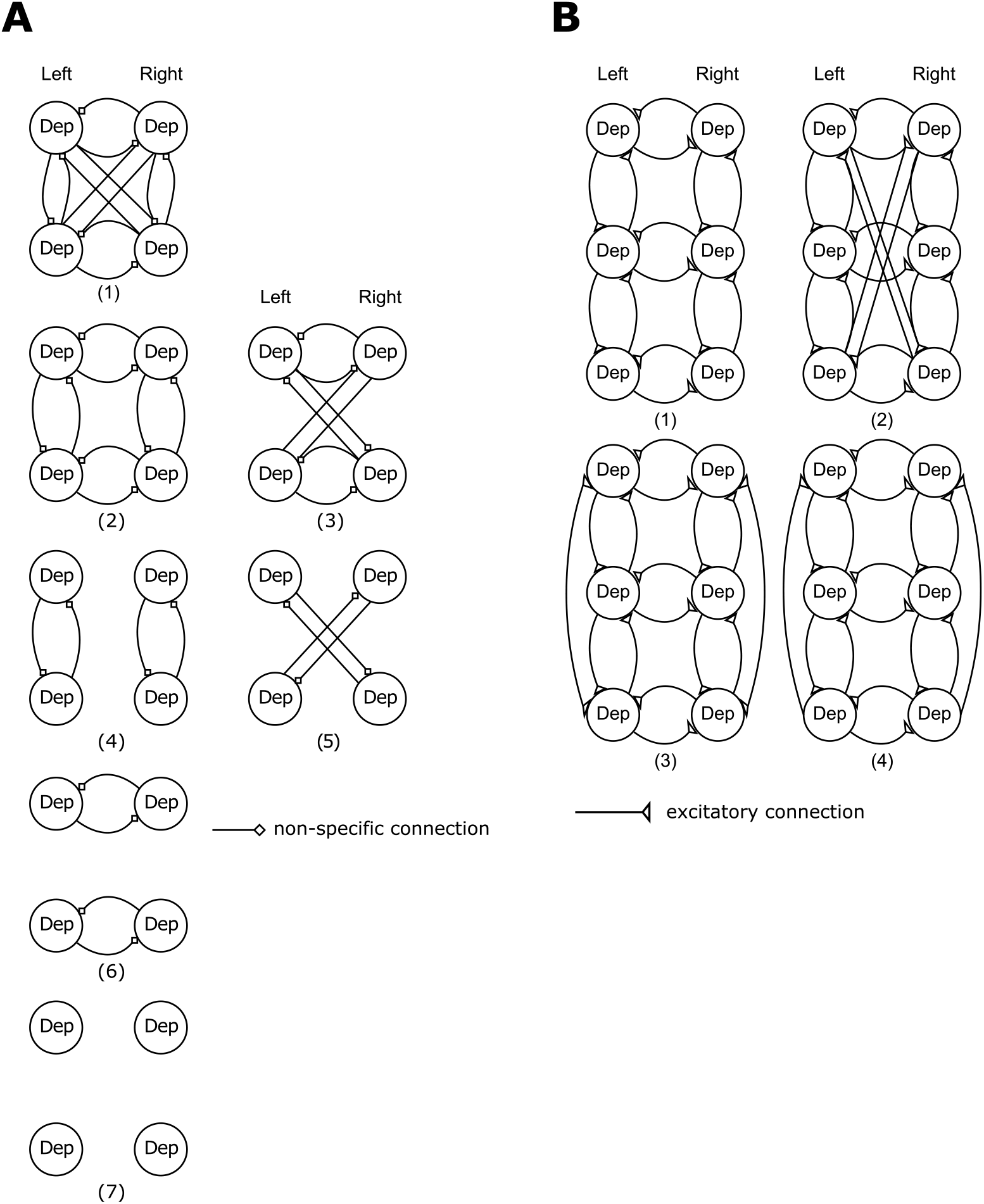
Model space for BMS. A: Model architectures tested with DCM for meso-meta and pro-meso recordings. B: Model architectures tested with DCM for pro-meso-meta recordings.

Significant differences in connectivity strengths were in both approaches (DCM and PC) determined by means of t-tests. In the DCM approach we assumed left-right symmetry and thus assigned both directions of a connection between two CPGs to the same connection (e.g. connection strengths from the left mesothoracic to the right mesothoracic and from the right mesothoracic to the left mesothoracic ganglion were averaged). This does not apply to the PC approach, since no directionality of the connections can be obtained there.

## 2 Results

In this study, we report results obtained with a method commonly used to analyze M/EEG data. We applied it to analyze the coupling strengths of pharmacologically induced rhythmic MN activity of the stick insect. Coupling between the motor systems of the CTr-joints of several segments, such as meso-meta, pro-meso was investigated; first, mathematical models that reproduce characteristic properties of the recorded depressor MN activity have been created with DCM which were then validated by means of the phase relations of the bursting activities (PC approach) and by an experimental condition with known connectivity, i.e. cut connectives between segments. After this careful validation, DCM was used to predict the coupling structure of the whole stick insect walking system involving all three (pro-, meso-, meta-) thoracic segments.

### Meso-Meta thoracic ganglia

In the first part of the experiments, the activity of the contralateral C2 nerves of the meso- and metathoracic ganglia was recorded. Data from both sides in the meso- and metathoracic ganglion were collected in 10 animals.

We used these bilaterally recorded data for the DCM analysis. First, we tested a number of possible predefined coupling structures. The models tested consisted of fully connected ganglia (1), circularly connected ganglia (2), cross connected ganglia with (3) and without intrasegmental connections (5), intrasegmentally unconnected (4), intersegmentally unconnected ganglia (6) and fully unconnected ganglia (7) (Fig 3 A).

All models were tested with excitatory and inhibitory connections. Here, we show the results with excitatory coupling, only, since in both cases (excitatory and inhibitory), the same winning model structure emerged but the best fitting model with excitatory connections had a higher probability according to the Bayesian Model Selection (BMS) procedure. BMS also showed that model (2) with the circular coupling structure best fitted the recorded data (Fig 4 A), since log-evidence and model posterior probability were highest.

**Fig 4.**
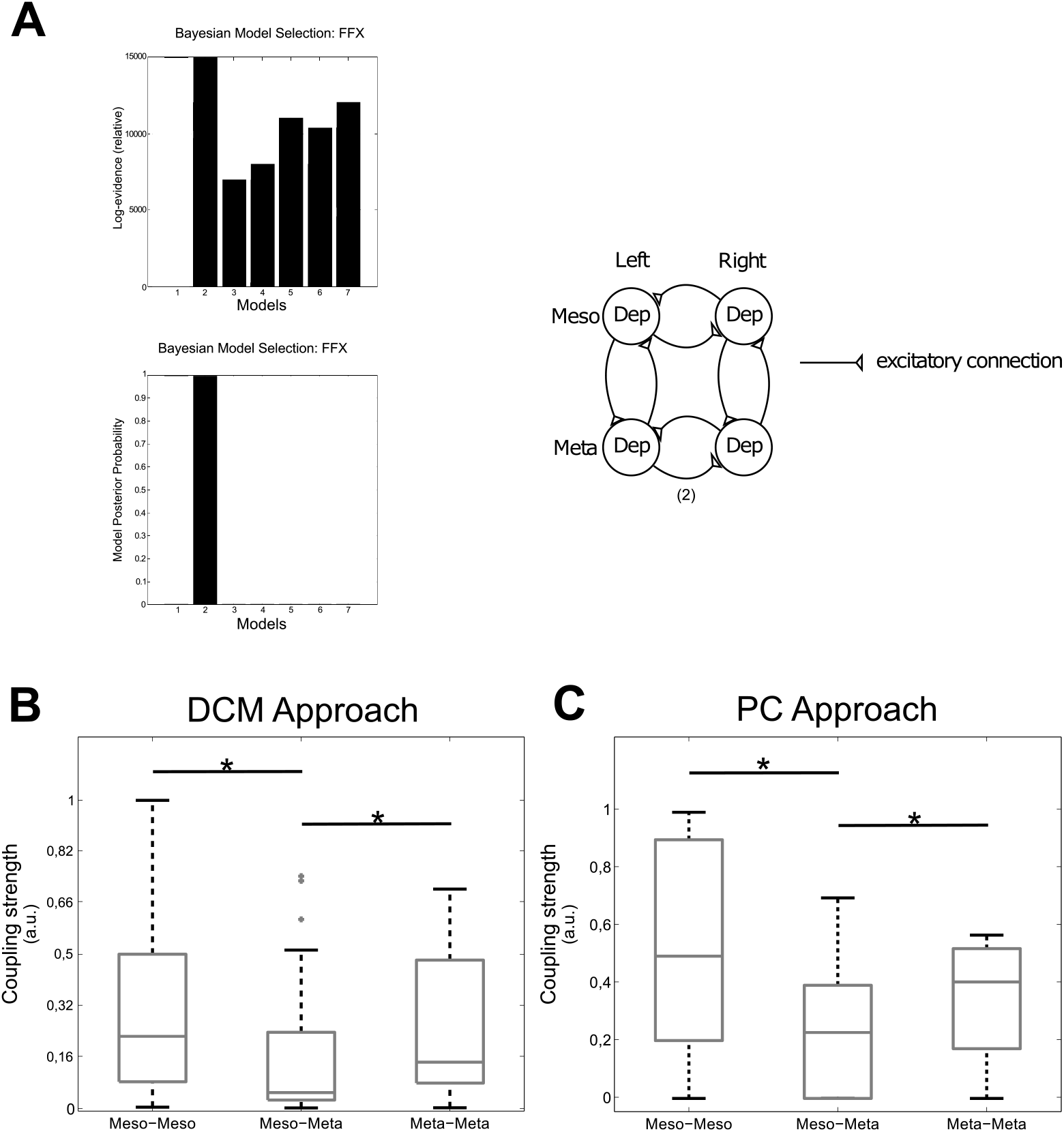
Connectivity meso-meta. A: BMS results for meso-meta DCM (left) and winning model structure (right). B: Boxplot showing the connectivity strengths for the DCM approach. The connectivity strengths were calculated for intervals that were pre-identified as phase-coupled by the PC approach. C: Boxplot showing the connectivity strengths for the PC approach. Meso-Meso denotes coupling between both sides of the mesothoracic and meta-meta the one between both sides of the metathoracic ganglion. Meso-Meta represents the intersegmental coupling between the meso- and the metathoracic ganglia. * denotes statistical significant differences.

We used this coupling structure as the basis for the estimation of the coupling strengths.

As first step of validation of DCM, we applied the DCM approach to *biased* intervals of the recorded data, i.e. to intervals that were pre-identified as phase-coupled in the PC analysis. We did this, since we were interested in the coupling strengths during coupled rather than uncoupled epochs.

The DCM analysis of the *biased* (phase synchronized) intervals yielded the coupling strengths, which are depicted in Fig 4 B. For a better comparison of the results we normalized the maximal coupling strength obtained from DCM to 1 (a.u.). We found the strongest coupling in the meso-meso and meta-meta connectivity, while the meso-meta connectivity was significantly weaker (tested with a two-sample t-test compared to meso-meso: *p* = 0.001, and compared to meta-meta: *p* = 0.044)(Fig 4 B).

The results of the phase-coupling (PC) approach are presented in Fig 4 C. In agreement with the DCM results, the likelihood was highest for the intrasegmental couplings, i.e. the coupling between both sides of the mesothoracic ganglion (meso-meso) and the coupling between both sides of the metathoracic ganglion (meta-meta). A two-sample t-test showed that the likelihood of intersegmental coupling (meso-meta) was significantly lower (*p* = 0.0396) than the intrasegmental couplings, while comparison of the intrasegmental couplings of meso-meso and meta-meta types showed no significant difference (*p* = 0.2224) between them.

Following, we performed a second validation to test whether DCM, which was developed for human brain data, can reasonably be applied to extracellular nerve recordings from animals. Therefore, we conducted an additional experiment where the connectives between the meso- and metathoracic ganglia were cut during the recordings. This surgical manipulation completely destroyed the connectivity between the two segments. We originally set up DCM to calculate coupling strengths of the fully connected system (A-matrix). Changes from this (cut connectives) were taken into account in the B-matrix (cf. Eq. (1)). We did not provide any prior information on which connections should be changed by DCM. The model showed a strong decrease of the connection strength, 90% on the left side and 95% on the right side, between the meso- and metathoracic ganglion (Fig 5). The connection between both segments was not completely removed by DCM. This is due to the fact that DCM is constructed to use a minimal connection strength whenever it is assumed to be present. In addition to the intersegmental decrease, there was a strong increase in the intrasegmental coupling in the mesothoracic ganglion (by factors of 2-20) and a decrease in connectivity in the metathoracic ganglion (by a factor of 2). This is in agreement with [28] where the authors could demonstrate that the mesothoracic ganglion showed intrasegmental phase-coupling even in the isolated state, while the connection of the metathoracic ganglion to the mesothoracic ganglion had to be present in order to detect robust in-phase coupling in the metathoracic ganglion.

**Fig 5.**
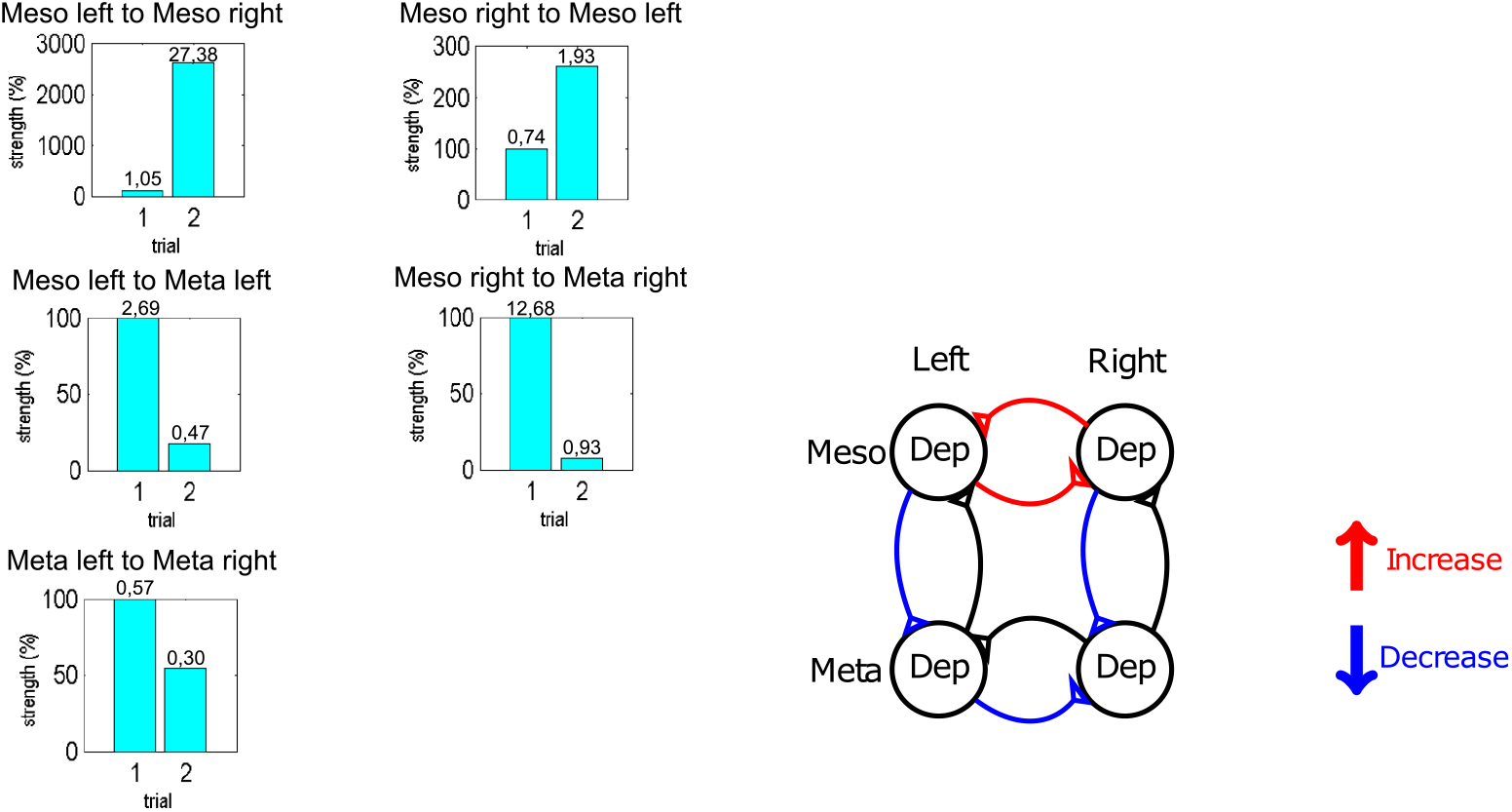
Second validation of DCM. Significant differences in connectivity from meso-meta thoracic connected segments (trial 1) to unconnected segments (trial 2). Left: Changes of connectivity strength, with trial 1 normalized to 100% coupling strength, absolute values of coupling strengths can be found on top of each bar; Right: Changes of connectivity compared to trial 1, red lines mark an increase in connectivity, blue lines a decrease in connectivity and black lines no reliable connectivity change (above 70%).

### Pro-Meso thoracic ganglia

To investigate the coupling between the pro- and the mesothoracic ganglia we recorded the activity of the C2 nerve on both sides of both ganglia in 5 animals, on both sides of the mesothoracic ganglion and on one side of the prothoracic ganglion in 3 animals and on both sides of the prothoracic and on one side of the mesothoracic ganglion in 5 animals.

We then analyzed these data with the DCM approach and tested the same model architectures as for the meso-meta recordings (see Fig 3, A). The BMS revealed the same result as for the meso-meta thoracic ganglia (Fig 6, A). That is why we used the same coupling structure for the analysis of the coupling strengths in the pro-meso thoracic ganglia (cf. Fig 4, A right). Again, we used the PC approach to *a priori* detect *biased*, i.e. phase-coupled intervals. Applying the DCM approach to these intervals, we can see that the intrasegmental meso-meso coupling (*p* = 0.0084) and the intersegmental pro-meso coupling (*p* = 0.0092) are significantly stronger than the intrasegmental pro-pro coupling as shown by two-sample t-tests (Fig 6, B).

**Fig 6.**
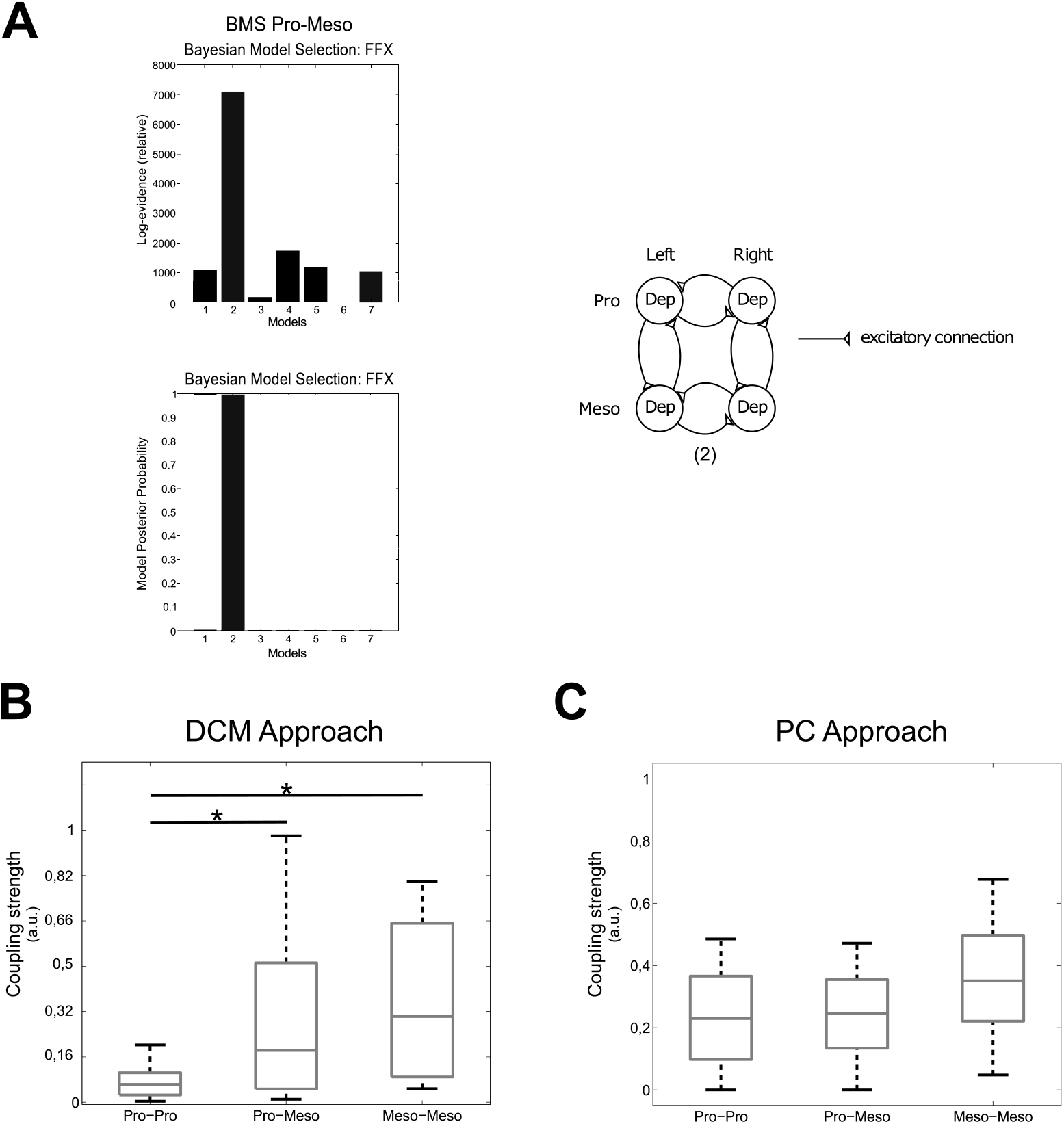
Connectivity pro-meso. A: BMS results for pro-meso DCM (left) and winning model structure (right). B: Boxplot showing the connectivity strengths for the DCM approach. The connectivity strengths were calculated for intervals that were pre-identified as phase-coupled by the PC approach. C: Boxplot showing the connectivity strengths for the PC approach. Pro-Pro denotes coupling between both sides of the prothoracic and meso-meso the one between the both sides of the mesothoracic ganglion. Pro-Meso represents the intersegmental coupling between the pro- and the mesothoracic ganglia. * denotes statistical significant differences.

We observed long intervals of tonic SDTr activity during the recordings of the connected pro- and mesothoracic ganglia, i.e. the small units in Fig 1 B, which were not present during the meso-meta recordings. This led to the detection of additional burst onsets by the previously described Poincaré section. Therefore, we now used a modified Poincare section defined by a k-means cluster (see Materials and methods) in order to filter out the bursts produced by those units. This basically amounted to applying a higher amplitude threshold to the data (cf. Fig 2, A right; adjustment of Poincaré section). Statistical analysis of the PC approach of the adjusted phase-differences using two-sample t-tests revealed no significant differences between all couplings (pp-mm *p* = 0.29, pm-mm *p* = 0.541, pm-pp *p* = 0.437) (Fig 6, C).

### Summary of DCM validation

We used the above as validation of the DCM results. For this, we compared the *relation* of coupling strengths obtained from the PC approach, which represent the ratio of phase-coupled intervals to total recording time, with the ones received by DCM analysis of *biased* intervals, as determined by the PC approach, in meso-meta and pro-meso recordings. In meso- and metathoracic recordings both approaches showed similar relations between analyzed ganglia, i.e. significantly stronger meso-meso and meta-meta compared to meso-meta connectivity (cf. Fig 4 B and C). In pro- and mesothoracic recordings we found a significantly decreased coupling strength in the pro-pro connection compared to all other connections using the DCM approach. This relation was not found by the PC approach, which might be due to a lowered signal-to-noise ratio (cf. Fig. 1) in this experimental setup (see Discussion). Additionally, we validated the DCM approach by introducing an experimental condition with prior knowledge on the connectivity, i.e. we cut the connectives between the meso- and metathoracic ganglia and thus effectively removed this connection. DCM was capable of portraying this condition by modulating the coupling strengths, i.e. by drastically reducing the connectivity strength of the experimentally removed connections.

### Pro-Meso-Meta thoracic ganglia

For the analysis of coupling between the pro-, meso- and metathoracic ganglia, we recorded the activity of the C2 nerve on both sides of all three ganglia in 3 animals. As the prior validation steps have shown a good agreement of the results of an established connectivity measure (PC approach) with the novel DCM approach, we applied DCM to these data to predict the overall network architecture and its coupling strengths of the thoracic nerve cord of the stick insect without *a priori* defining a proxy for phase-coupled intervals. We used a gliding window of 50 s duration to select five intervals for each animal instead. The models tested with BMS consisted of a union of the previously discussed networks (1), with added cross connections (2), as well as bi- (3) and unidirectional lateral connections (4) between pro- and metathoracic ganglia (Fig 3 B). Model 4 became the winning model (Fig 7, A). Thus, the BMS suggests that the full network consists of the union of the subnetworks and a lateral feedback from meta- to prothoracic ganglia (Fig 7 B). An a posteriori t-test on the coupling strengths revealed no significant difference between the nine connections (all *p*’s > 0.05). A further t-test on the meso-meso coupling strengths in meso-meta and pro-meso-meta recordings revealed a significantly reduced connectivity in the latter recordings (*p* = 0.0044).

**Fig 7.**
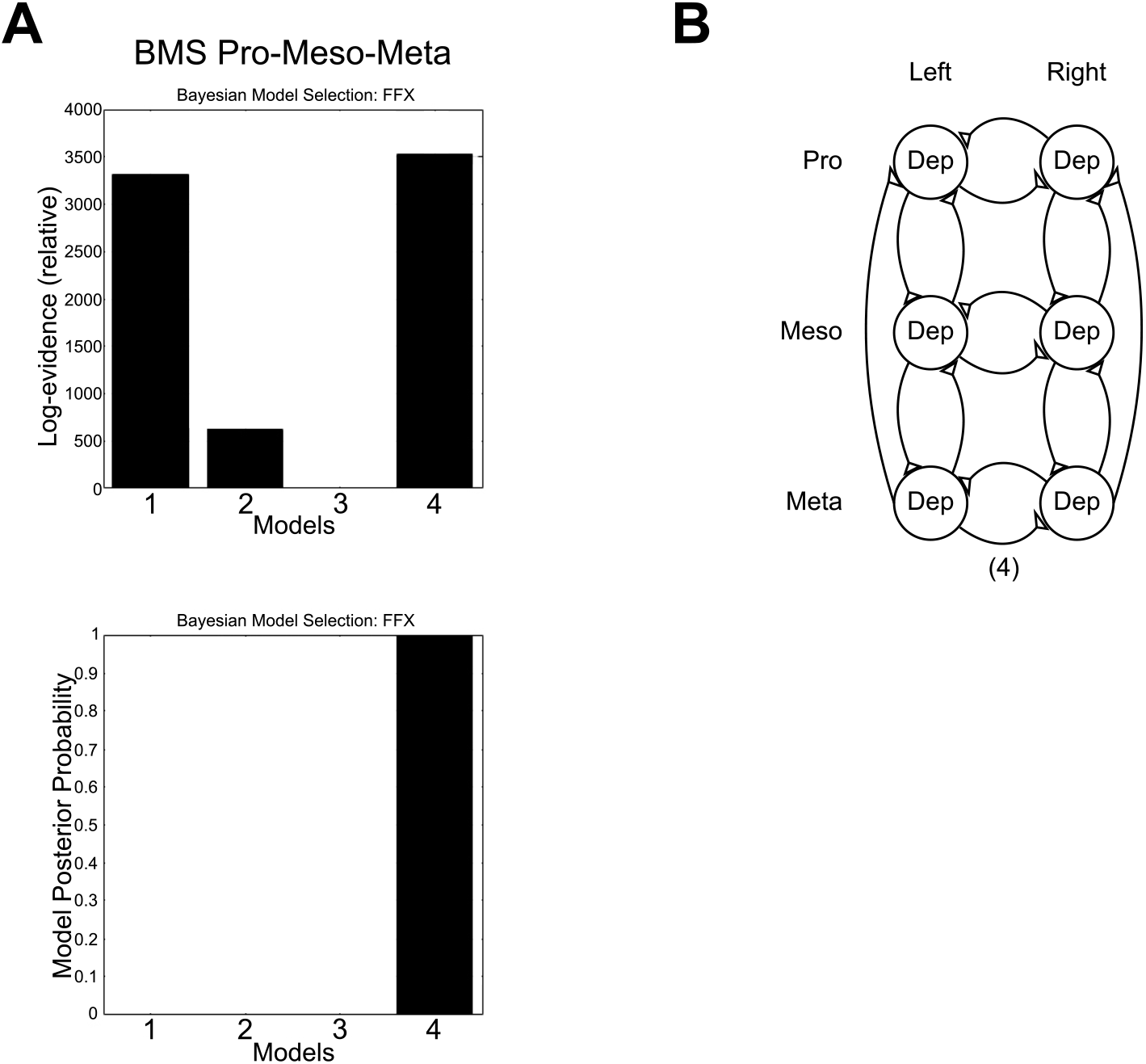
Network structure: BMS pro-meso-meta. A: BMS results: Log evidence (top) and posterior probability (bottom) for pro-meso-meta recordings are highest for model (4); B: Representation of the winning network structure model (4). Triangles denote excitatory connections.

## 3 Discussion

### General coupling structure of CPG networks

In this paper, we, for the first time, provide results on the coupling structure and the coupling strengths for the whole thoracic nerve cord in a multi-legged, deafferented locomotor system. Our results provide strong evidence that in all animals investigated, the inter- and intrasegmental central couplings follow the same organizational principle. In Fig 8, we sum up the structure and the strengths of the network connectivity from the different experimental setups. BMS on the set of various fitted DCM models preferred an eight-shaped structure (see Fig. 7 B) consisting of ipsilateral inter- and contralateral intrasegmental connections of the CPGs and thus predicted a high probability for the existence of these connections. Importantly, the models containing cross connections, e.g. from pro-left to meso-right, models with missing inter- or intrasegmental coupling and fully connected models were not selected by BMS. The coupling structure we propose here is in accordance with previous results showing that ipsilateral leg coordinating influences are indeed transmitted by the ipsilateral connective [8, 13, 24]. We restricted the tested model architectures to those containing all excitatory or all inhibitory connections without any mixture of both types to limit the model space for this first application of DCM.

**Fig 8.**
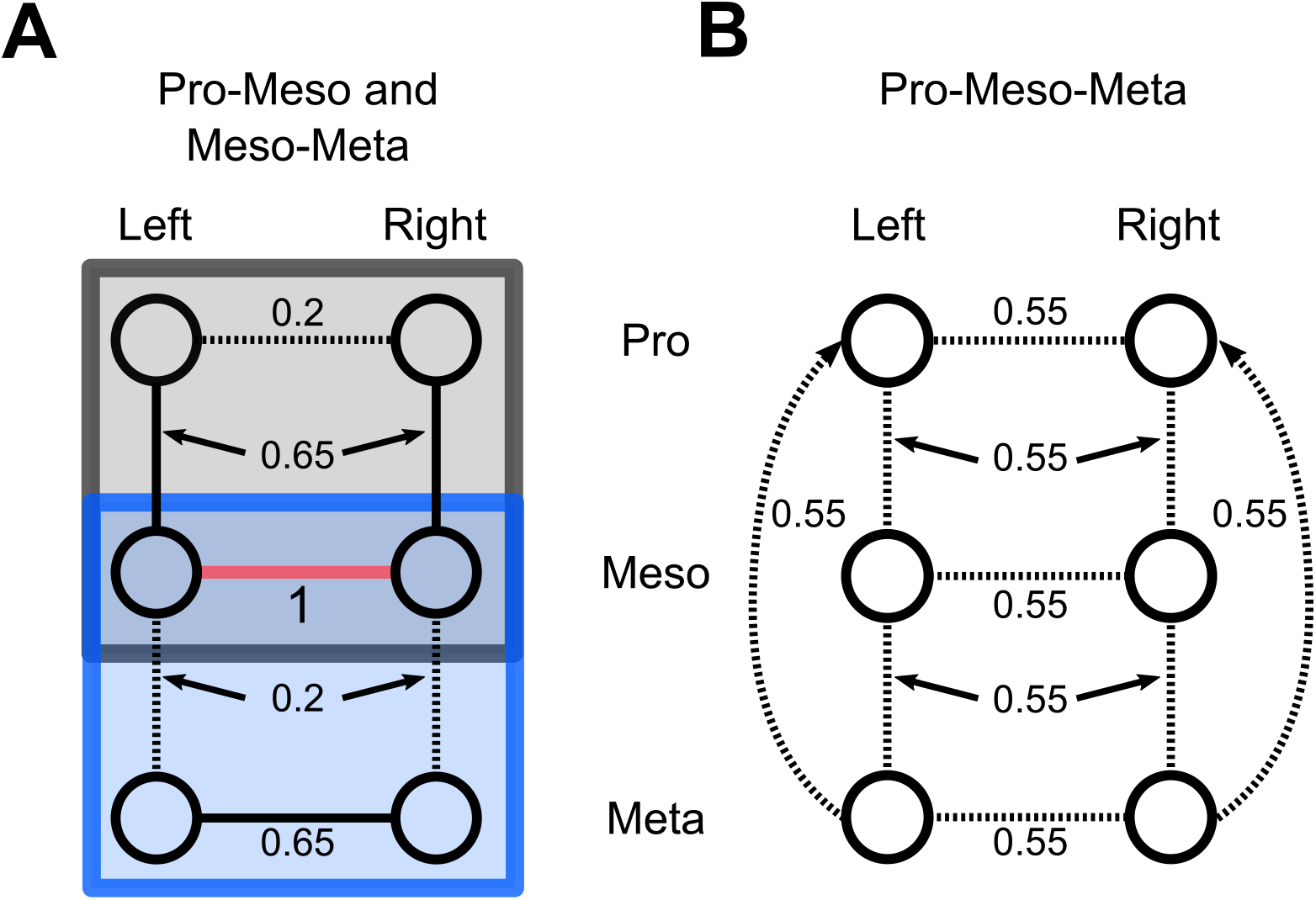
Summary of the coupling strengths (numbers) obtained by the DCM approach. A: Recordings involving two ganglia: pro-meso recordings marked by a grey rectangle and meso-meta recordings with a blue rectangle, respectively. B: Recordings involving pro-, meso- and metathoracic ganglia. All coupling strengths are normalized to the meso-meso connectivity (marked with magenta line) strength in the corresponding meso-meta recordings (blue rectangle). Significantly smaller connectivity strengths are represented by dashed lines.

The analysis of coupling strengths in the pro-meso-meta recordings showed only minor insignificant differences (identical up to the third decimal place) between the various connections (Fig. 8 B). This might have several reasons. The recording of all three pairs of ganglia might have introduced noise into the system which reduced the coupling strengths detected in the meso-meta and pro-meso recordings. Additionally, since we used a gliding window approach instead of biased intervals, there might also be fluctuations in coupling strengths which were averaged out over time. Further, the meso-meso coupling, which we used as a normalized reference, was significantly reduced compared to the pro-meso and meso-meta recordings. This might reflect the stick insects flexibility in coupling and decoupling of certain legs into the lcocomotor system to produce different movement patterns or behaviours, e.g. searching movements.

### Validation and discussion of the DCM approach

We used three steps of validation of the DCM approach. In a first step, we evaluated the connectivity with a well established phase-connectivity approach in addition to DCM for the meso- and metathoracic recordings. Both approaches showed similar results, i.e. significantly stronger meso-meso and meta-meta compared to meso-meta connectivity (cf. Fig 4 B and C). In a second step, we introduced an experimental condition with prior knowledge on the connectivity, i.e. we cut the connectives between the meso- and metathoracic ganglia and thus removed this connection. Here, we could show that the DCM approach was capable of reproducing “known” coupling strengths in the system. When unspecified changes of connectivity were entered into the model, this resulted in a drastic reduction in intersegmental connectivity strength. As we could not remove any connection from the model without biasing DCM, it still assigned some connectivity strength to the intersegmental connections even though they were not present in the animal. In a third step, we again compared the results from the DCM and PC approaches for the pro- and mesothoracic recodings. While we found a significantly decreased coupling strength in the pro-pro connection as compared to all other connections in the DCM approach, this difference was not significant in the PC approach (cf. Fig. 6 B, C). Adding the prothoracic ganglion to the network to be tested lowered the signal-to-noise ratio (cf. Fig. 1), which might have a bigger effect here, since the exact determination of burst onsets is crucial for the PC approach.

Since the majority of validations revealed a good agreement between results, we are confident to state that even though DCM was developed for neural networks in the human brain, it is appropriate to be used for less complex systems, i.e. insects thoracic nerve cords.

The application of DCM for the analysis of central intra- and intersegmental CPG interactions has several advantages, but also some disadvantages. In contrast to methods like coherence, cross-spectra, phase-coupling or phase-delay analyses, which give an incomplete picture for stationary signals and cannot be interpreted in terms of directionalities [16], DCM, by modeling the system causing the observed effects, is able to give insight into the effective connectivity strengths, as well as the global coupling architecture of the studied system. Additionally, DCM is able to deal with fewer experimental recordings, since it is used to estimate the connectivity of subintervals (e.g. 50 s epochs in this article) and is therefore still capable of reconstructing the network structure from this data. A disadvantage for the application of DCM to weakly coupled locomotor systems, e.g. in the stick insect, is that it requires a steady state like behavior. While DCM is still able to predict the overall network structure, it seems not to be able to detect differences in coupling strengths between the investigated connections if steady-state behavior is not guaranteed (see preceeding paragraph). Thus prior detection of such steady states by other measures (e.g. PC approach) seems to be necessary for a meaningful interpretation of the obtained connectivity strengths. In other locomotor systems with stronger central coupling, e.g. cockroach, we anticipate this disadvantage to be less pronounced as the connectivity strength is expected to be more stable over recordings and thus relies less on the selection of epochs for estimation.

### Central intersegmental CPG interactions in other locomotor systems

Central intersegmental CPG interactions have been demonstrated in the past using various animal models. In-phase intersegmental activity of MN pools has been reported earlier for deafferented preparations of the crayfish [41] and the stick insect [5] after pharmacological CPG activation. In the deafferented thoracic ganglia of the locust, intersegmental depressor MN activity also expressed similar behavior [24]. In contrast, data of [38] had suggested ipsilateral coupling between ipsilateral levator and depressor MN activity in adjacent ganglia of the locust. In the deafferented thoracic nerve cord of the hawk moth, pharmacological activation of the depressor MN pools produced an activity pattern that resembled the tripod leg coordination pattern that emerges during walking in a large number of insect species [22]. Finally, similar intersegmental coordination patterns were recorded in the interconnected meso- and metathoracic ganglia of the cockroach thoracic nerve cord with the sub-esophageal ganglion (SEG) attached to it [12, 17]. Thus, centrally-generated motor patterns in all the above mentioned preparations revealed intersegmental coupling of activity among CPGs.

### Stabilizing effect of the mesothoracic ganglion

Our results suggest that the intrasegmental coupling of the mesothoracic ganglion is stronger than other connections (Fig 8, cf. Figs 4 B, 6 B). Cutting the connectives between the meso- and metathoracic ganglia, showed that the presence of the ipsilateral intersegmental meso-meta connection was needed for a strong intrasegmental coupling in the metathoracic ganglion. Our result supports the findings by [28] where the authors showed that phase-coupling of neural activity in the metathoracic ganglion is more stable when the meso- and metathoracic ganglia are interconnected. Here, we found an overall increased meso-meso connectivity after cutting the connectives between the meso- and metathoracic ganglia, while Mantziaris and colleagues [28] observed a slight increase in regularity in meso-meso phase distributions in favor of in phase activity, i.e. a decrease in variability of interburst phase differences, when both (meso- and metathoracic) ganglia were connected. This might be due to the fact, that the DCM approach does not distinguish between in-phase and out-of phase synchronization of the mesothoracic ganglion. Our results suggest that a higher meso-meso coupling is necessary to model the remaining rhythmic behavior after the loss of metathoracic inputs. In a behavioral study Grabowska et al. could show that stick insects with amputated middle legs display a malfunction of coordination with multiple stepping of front and hind legs as well as ipsilateral legs being in swing phase simultaneously [18]. Combining these findings, we suggest that the activity of the mesothoracic CPGs stabilizes that of the metathoracic segments, ensuring a stable rhythm in the meso- and metathoracic ganglia.

### Weak coupling in the prothoracic ganglion

The intrasegmental coupling strength of the prothoracic ganglion was the weakest (Fig 8 A). In the analysis of the pro-meso-meta ganglia no significant differences between the coupling strengths within the network could be seen, while the reference connection (meso-meso) was significantly reduced (in average to 55%) compared to that in meso-meta recordings (Fig 8 B). Moreover, when adding the prothoracic ganglion to the network to be analyzed (pro-meso and pro-meso-meta activity), the signal-to-noise ratio decreased, reducing the aforementioned stabilizing effect of the mesothoracic ganglion. These results might hint at the special role the front legs have. It has been shown that the front legs can perform additional steps or searching movements independently of other legs [7, 18]. Our results suggest that this might be achieved by a weaker lateral intrasegmental coupling between the prothoracic CPGs and a weaker ipsilateral intersegmental coupling between pro- and mesothoracic CPGs. In the deafferented stick insect preparation, restricted CPG activation in the prothoracic ganglion had indeed no intersegmental effect on the mesothoracic networks [25]. In contrast to this, a recent study by Knebel et al. has reported a strong in phase coupling of pro- and mesothoracic ganglia after restricted activation, i.e. using a split bath preparation, of the prothoracic ganglia in the locust [24]. Furthermore, it has been shown in cockroaches that intrasegmental, i.e. meso-meso activity has a strong anti-phase relationship in the absence of sensory feedback [17]. This is a requirement for producing the tripod coordination pattern, which is preferred by these animals. This behavior is further enhanced by sensory feedback from a single stepping front leg, suggesting an additional stabilizing effect by the prothoracic ganglia. These findings suggest a stronger coupling between pro- and mesothoracic ganglia and less flexibility in front leg movements in locusts and cockroaches.

### Coupling from meta-to prothoracic ganglion

The model by Daun-Gruhn & Tóth hypothesizes that the existence of a connection from the hind to the front leg is necessary to establish coordinated locomotor patterns in the three ipsilateral legs [11]. The recordings from all three ganglia enabled us to gain insights into the coupling structure involving the pro- and metathoracic ganglia. Our DCM results support the hypothesis of Daun-Gruhn & Tóth by revealing an ipsilateral feedback from the meta-to the prothoracic ganglia without any contralateral connection. Knebel et al. (2017) could show a clear coupling of pro- and metathoracic ganglia in the case of restricted activation of prothoracic as well as restricted activation of metathoracic ganglia [24]. In contrast to the results presented here, this finding suggests the presence of a bidirectional coupling of both ganglia in locusts.

Furthermore, we found no significant differences in coupling strengths in the case of simultaneous recordings from all three ganglia, while it had previously been shown that depressor activity in all segments seems to be weakly phase coupled in locusts [24].

### Comparison with a connectivity model of leg coordination in the cockroach

David and colleagues (2016) reported CPG coupling strength in the meso- and metathoracic ganglia of the cockroach by calculating the transition latencies and phase relations between bursts of activity [12]. In contrast to the results presented here for the stick insect, ipsilateral connections in the cockroach system were found to be stronger than the contralateral ones, while diagonal coupling interactions were also present in the resulting connectivity scheme reported. Moreover the meso-to metathoracic, descending coupling was weaker than the ascending. Such an asymmetry was not systematically observed in our results. In addition, as reported in [12], intrasegmental metathoracic coupling was stronger than coupling between the mesothoracic hemisegments, whereas the opposite is demonstrated here for the stick insect.

The reasons for these discrepancies between the cockroach and the stick insect systems are not well understood. Importantly, the subesophageal ganglion (SOG) and the abdominal ganglia in the aforementioned study were left attached to the thoracic ganglia. Thus, descending or ascending signals, or both, might affect CPG coordination and coupling strength. Our experiments were performed in the absence of SOG input and we should therefore exercise care when comparing our results to those of David et al. [12].

In a recent modelling study, Szczecinski et al. [42] demonstrated that interleg coordination patterns in *D. melanogaster* result from the interplay between static stability of the animal and robustness of the coordination pattern. The authors found that at a large variety of walking speeds (hence coordination patterns), only the ipsilateral phase differences change, whereas contralateral phase differences remain at about 1/2. Their simulation results support this finding. Taking this result into consideration, a stronger ipsilateral coupling can be expected in cockroaches, since they usually exhibit tripod coordination pattern during walking. By contrast, the slow walking stick insects may require weaker ipsilateral coupling that could be affected more strongly by afferent sensory signals. To our knowledge, there is no similar study concerning static stability in other insects. However, it may be that ipsilateral phase relationships are critical for intersegmental leg coordination in other insects as well. It would be interesting to know whether the differences in intra- and intersegmental coupling between stick insects and cockroaches are related to the variable static stability of the two animals in relation to their inherent walking speed.

## 4 Conclusions

In this paper, we have shown for the first time that well-established methods for analyzing M/EEG data can be adapted for the analysis of pilocarpine-induced fictive locomotor patterns of the stick insect *Carausius morosus* to estimate coupling architecture and strengths from real data. Applying the DCM approach, we could predict a high probability for the existence of ipsilateral inter- and lateral intrasegmental connections of the CPGs. A connection from the meta-to the prothoracic ganglion was also hypothesized. We could further discern changes in the connectivity of the thoracic ganglia of the stick insect in different experimental conditions. DCM detected the absence of coupling between the meso- and metathoracic ganglia after the connectives had been cut. Using DCM, we have established that the intrasegmental mesothoracic connectivity is the strongest from all others in all three thoracic ganglia. Moreover, this coupling has a stabilizing effect on intrasegmental metathoracic activity. Connectivity involving prothoracic ganglia, by contrast, is either weak or depends on the specific network topology to be analyzed. This could account for the fact that the prothoracic ganglia have to allow decoupling of the front legs from the rest of the locomotor system to enable search movements of the front legs independently of the middle and the hind leg movements.

## Acknowledgements

We thank Anke Borgmann for useful discussions on the analysis. This study was funded by the University of Cologne Emerging Groups Initiative (CONNECT) within the framework of the Institutional Strategy of the University of Cologne and the German Excellence Initiative. SD gratefully acknowledges additional support from the German Research Foundation, Germany (DA1953/5-2). CM was an associate member of the RTG 1960 ”Neural Circuit Analysis on the Cellular and Subcellular Level” funded by the DFG.

